# C-di-AMP levels modulate *Staphylococcus aureus* cell wall thickness as well as virulence and contribute to antibiotic resistance and tolerance

**DOI:** 10.1101/2023.04.18.537236

**Authors:** Vanina Dengler Haunreiter, Andrea Tarnutzer, Julian Bär, Manuela von Matt, Sanne Hertegonne, Federica Andreoni, Clément Vulin, Lisa Künzi, Carmen Menzi, Patrick Kiefer, Philipp Christen, Julia A. Vorholt, Annelies S. Zinkernagel

**Author notes:** Address correspondence to Annelies S. Zinkernagel. Vanina Dengler Haunreiter and Andrea Tarnutzer contributed equally. Order was determined by seniority.

## Abstract

Beta-lactam antibiotics are widely used to treat infections caused by the important human pathogen *Staphylococcus aureus*. Resistance to beta-lactams, as found in methicillin-resistant *S. aureus* (MRSA), renders effective treatment difficult. The second messenger cyclic di-3′,5′- adenosine monophosphate (c-di-AMP) promotes beta-lactam resistance in clinical *S. aureus* isolates. C-di-AMP plays a crucial role in the regulation of cellular processes such as virulence, cell wall homeostasis and resistance to beta-lactams in many bacterial species. In *S. aureus,* c-di-AMP synthesis is mediated by the diadenylate cyclase DacA, while its degradation is carried out by the phosphodiesterases GdpP and Pde2.

In this work, we assessed the effect of altered c-di-AMP levels due to mutations in *cacA*, *gdpP* or *gdpP/pde2* on virulence determinants. We report that a previously described growth defect in bacteria producing high c-di-AMP levels is mainly attributable to smaller cell size. High c-di-AMP levels also led to decreased survival upon oxidative stress, reduced production of the antioxidant staphyloxanthin, increased oxacillin and fosfomycin resistance and increased cell wall thickness. While resistance to ceftaroline was not affected, high c-di-AMP levels promoted tolerance to this antibiotic. In response to cell wall stress induced by antibiotics, the three-component regulatory system VraTSR mediates an increase in cell wall synthesis via the cell wall stress stimulon (CWSS). Increased c-di-AMP levels led to an activation of the CWSS. Upon deletion of *vraR*, resistance to oxacillin and fosfomycin as well as cell wall thickness diminished in the Δ*gdpP* mutant, indicating a contribution of the VraTSR system to the cell wall related phenotypes.

**Importance:** Antibiotic resistance and tolerance are substantial health-care related problems, hampering effective treatment of bacterial infections. Mutations in the phosphodiesterase GdpP, which degrades cyclic di-3′, 5′-adenosine monophosphate (c-di-AMP), have recently been associated with resistance to beta-lactam antibiotics in clinical *Staphylococcus aureus* isolates. In this study, we show that high c-di-AMP levels decreased the cell size and increased the cell wall thickness in *S. aureus* mutant strains. As a consequence, an increase in resistance to cell wall targeting antibiotics such as oxacillin and fosfomycin as well as in tolerance to ceftaroline, a cephalosporine used to treat MRSA infections, were observed. These findings underline the importance of investigating the role of c-di-AMP in the development of tolerance and resistance to antibiotics in order to optimize treatment in the clinical setting.

## Introduction

Antibiotic resistance and tolerance greatly hinder the effectiveness of bacterial infection treatment. Beta-lactam antibiotics are widely used to treat infections caused by the important human pathogen *Staphylococcus aureus*. However, resistance development as seen in methicillin-resistant *S. aureus* (MRSA) are commonly encountered in community- and hospital-acquired infections (1).

Second messenger signaling allows bacteria to rapidly respond to environmental changes including antibiotics. Cyclic di-3′,5′-adenosine monophosphate (c-di-AMP) is the most recently discovered dinucleotide second messenger and is present in at least 275 bacterial species, predominantly in the Gram-positive phyla Firmicutes and Actinobacteria, but also in some Gram-negative genera and in certain Archaea (2). In *S. aureus*, c-di-AMP is synthesized by the diadenylate cyclase DacA and degraded by two phosphodiesterases containing a DHH/DHHA1 domain, GdpP (membrane-bound) and Pde2 (cytosolic) (3, 4). GdpP degrades c-di-AMP to pApA in a single-step reaction while Pde2 converts c-di-AMP and pApA to AMP. In contrast to other species in which Pde2 plays the main role in c-di-AMP degradation, Bowman and colleagues found that, in *S. aureus*, Pde2 preferably degrades pApA to AMP while GdpP is mainly responsible for c-di-AMP degradation (4).

C-di-AMP is the only second messenger molecule known to be essential for growth under standard laboratory conditions (5). It is involved in various cellular processes, including virulence, salt and cell wall homeostasis, and resistance to beta-lactam antibiotics (6, 7). Recently, mutations in *gdpP* were shown to confer beta-lactam antibiotics resistance in clinical isolates lacking *mec* determinants and displaying normal PBP4 levels, emphasizing the clinical relevance of c-di-AMP levels in conferring resistance to antibiotics (8–10).

C-di-AMP is known to influence bacterial cell wall homeostasis; however, the underlying mechanisms are not fully understood and are likely species-specific. In *S. aureus*, high c-di-AMP levels result in increased peptidoglycan crosslinking and resistance to beta-lactam antibiotics while low c-di-AMP levels decrease beta-lactam antibiotics resistance (4, 11). Cell wall targeting antibiotics can activate the three-component regulatory system VraTSR, which regulates the expression of more than 40 genes belonging to the cell wall stress stimulon (CWSS), leading to increased cell wall synthesis and reduced expression of autolysins (12–14). A link between c-di-AMP levels and CWSS activation was found in two different *S. aureus dacA_G206S_* mutants where decreased c-di-AMP levels correlated with a decreased activation of the CWSS (11).

In this work, we investigated the effect of altered c-di-AMP levels on various *S. aureus* virulence determinants such as growth, oxidative stress response, cell wall thickness and antibiotic susceptibility as well as the connection with the CWSS.

## Results

### C-di-AMP indirectly influences growth characteristics through cell size

Previous studies have demonstrated in *S. aureus* that the absence of either one or both phosphodiesterases (Δ*gdpP*, *Δpde2* or Δ*gdpP/*Δ*pde2*) resulted in increased c-di-AMP levels and lowered the growth rate based on OD measurements (4, 15). In contrast, a *dacA_G206S_* mutant strain, producing lower amounts of c-di-AMP, grew comparably to the WT or slightly faster depending on the *S. aureus* strain background (4, 11, 16). Using the MRSA strain LAC* and its isogenic *dacA_G206S_*, Δ*gdpP* and Δ*gdpP/*Δ*pde2* mutants, we aimed to further dissect the impact of c-di-AMP levels on various growth characteristics. As expected, c-di-AMP levels were increased in the Δ*gdpP* mutant compared to the WT (supplementary table 1). The Δ*gdpP*/Δ*pde2* mutant showed even higher levels than the Δ*gdpP* single mutant while c-di-AMP levels decreased in the *dacA_G206S_* strain (Supplementary Table S1 and S2). Measuring optical density as a population size proxy, we observed an attenuated increase in OD in the mutants with higher c-di-AMP levels, as described before (4, 15) (Figure 1A). However, when measuring the colony forming units (CFUs) after growth in liquid TSB we detected a similar growth rate in all strains (Figure 1B). An effect due to viability can be excluded as we found no significant differences in cell viability measured by a live-dead stain with flow cytometry between WT, Δ*gdpP* and Δ*gdpP/*Δ*pde2* mutant strains (Supplementary Figure S1A).

**Figure 1:**
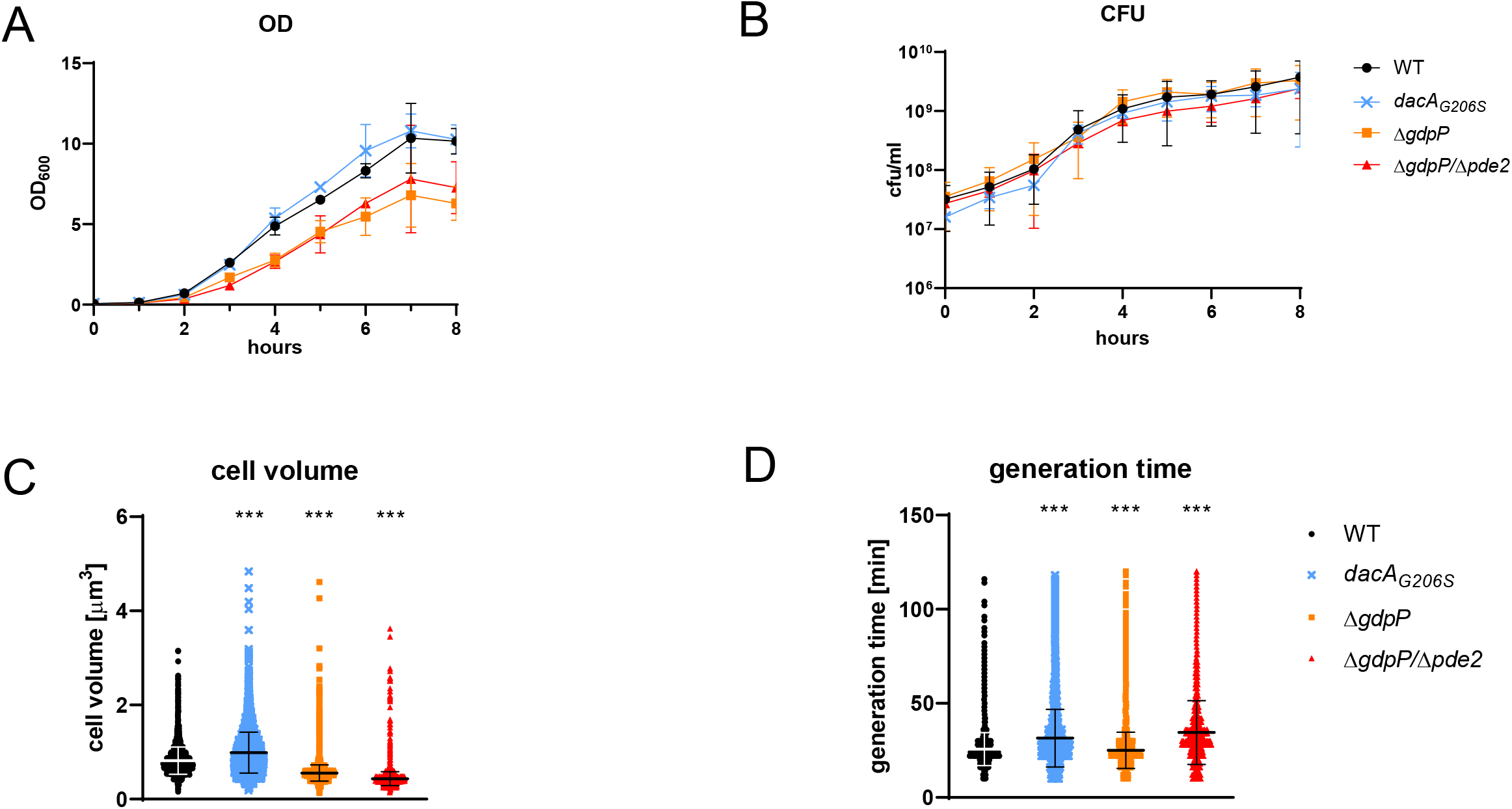
C-di-AMP indirectly influences growth characteristics through cell size. OD (A) and CFUs (B) were measured in TSB broth. Data displays mean ± SD of three biological replicates. Nonlinear regression showed no significant differences in CFUs. Cell volume (C) and generation time (time between division events) (D) of single cells was determined in a microfluidic microscope system. Each dot represents one cell and bars indicate mean ± SD (C, D). Statistical significance was assessed by a mixed effects model using a random intercept term for the biological replicate (three replicates) followed by estimated marginal means post-hoc tests comparing WT to each other strain including multivariate t distribution based p-value correction. *** p<0.001

It has been shown that c-di-AMP plays a role in controlling cell size (3, 16). As OD measurements depend on the cell size (17, 18), we assessed the volume of single cells growing in a mother-machine microfluidic device by single cell microscopy (representative movie of each strain in Supplementary Material). Compared to the WT (0.82 ± 0.29µm^3^), the mean cell volume significantly increased in the *dacA_G206S_* mutant (0.99 ± 0.43µm^3^, 134% of WT) and decreased in the Δ*gdpP* (0.55 ± 0.17µm^3^, 66% of WT) as well as in the Δ*gdpP/*Δ*pde2* (0.43 ± 0.15µm^3^, 50% of WT) mutant, suggesting that c-di-AMP affects cell volume in a concentration-dependent manner (Fig. 1C). In line with these results, the colony size measured 24 hours after inoculation on agar plates negatively correlated with c-di-AMP levels (Supplementary Figure S1B). Next, we assessed the generation time (defined as time between cell division events) and found no indication of impaired growth in the Δ*gdpP* mutant compared to the WT (25.03 ± 9.62 min and 25.68 ± 8.75 min, respectively) (Figure 1D). A prolonged generation time was found in both the *dacA_G206S_* (31.51 ± 15.27 min) and the Δ*gdpP/*Δ*pde2* (34.54 ± 16.92 min) mutants.

Based on these observations, we conclude that the reduction in OD-based growth in high c-di-AMP mutants is biased by the smaller cell size and does not reflect growth impairment in the sense of a reduction in cell division rate or viability.

### Elevated c-di-AMP levels lead to decreased survival under oxidative stress and reduced staphyloxanthin production

To assess how c-di-AMP levels affect *S. aureus* virulence, we quantified resistance to oxidative stress, a key virulence feature promoting survival in the host (19). Increased c-di-AMP levels led to a significantly lower survival after exposure to H_2_O_2_, whereas no statistical significance in survival was found in the *dacA_G206S_*mutant with decreased c-di-AMP levels (Figure 2A). Resistance to oxidative stress is linked to the virulence factor staphyloxanthin, an important antioxidant in *S. aureus* (20). Staphyloxanthin levels were therefore expected to be linked with survival after oxidative stress. Indeed, staphyloxanthin levels were significantly lower in strains with elevated c-di-AMP levels (Figure 2B).

**Figure 2:**
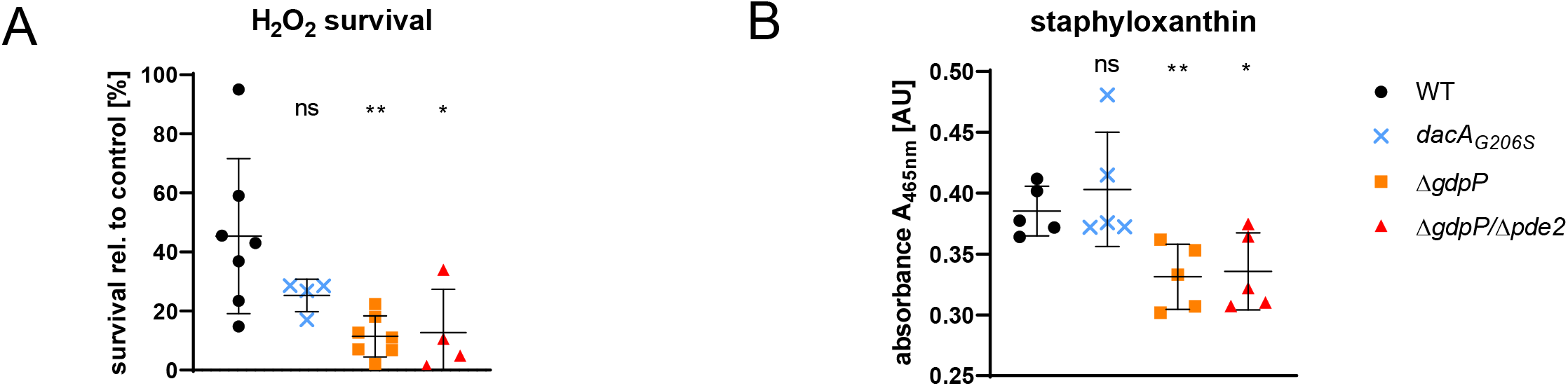
Elevated c-di-AMP levels lead to decreased survival under oxidative stress and reduced staphyloxanthin production. (A) Survival rate of stationary growth phase bacteria after one hour exposure to 30 mM H_2_O_2_ in PBS was calculated relative to the control (no H_2_O_2_). (B) Staphyloxanthin was quantified in cultures grown for 24 hours in TSB. Data display mean ± SD of at least 4 biological replicates, statistical significance was assessed by unpaired Student’s t-test comparing WT to each strain. *p < 0.05, **p < 0.01

### C-di-AMP modulates tolerance to ceftaroline and resistance to fosfomycin

The level of c-di-AMP positively correlates with resistance to beta-lactam antibiotics in many bacterial species (3, 21). Accordingly, the Δ*gdpP* and Δ*gdpP/*Δ*pde2* mutant strains displayed a 1000-fold increase in minimal inhibitory concentration (MIC) of the beta-lactam oxacillin (256 µg/ml) as compared to the WT strain (0.25 µg/ml) while the *dacA_G206S_* mutant was four-times more susceptible (0.064 µg/ml) (Table 1, upper part). Next, we evaluated susceptibility to ceftaroline, a fifth-generation cephalosporin displaying increased affinity to the mutated penicillin-binding protein 2a (PBP2a) of MRSA strains. We observed no changes in MIC in any of the mutants (Table 1, upper part). However, a time-kill curve with 40x MIC ceftaroline revealed differences in tolerance to ceftaroline between the strains (Figure 3A). After 1.5 and 3 hours growth in the presence of ceftaroline, the Δ*gdpP* and Δ*gdpP/*Δ*pde2* mutants survived significantly better than the WT. This difference became less pronounced after 6 hours and was undetectable after 24 hours. The *dacA_G206S_* strain showed a consistently lower survival rate than the WT and was completely eradicated after 6 hours of 40x MIC ceftaroline treatment. In summary, tolerance quantified by minimal duration of killing 99% of the population (MDK_99_) increased in the two mutants with high c-di-AMP levels and decreased in the *dacA_G206S_* mutant (Figure 3B).

**Figure 3:**
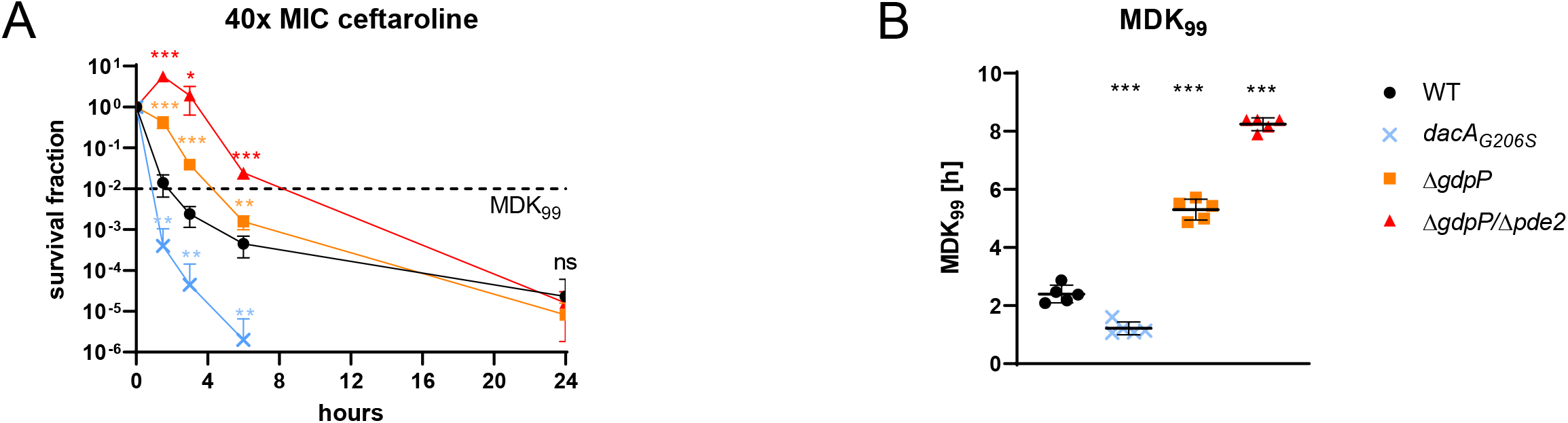
C-di-AMP modulates tolerance to ceftaroline. (A) The survival fraction of bacteria grown in TSB supplemented with 40x MIC ceftaroline was calculated relative to the inoculum. The horizontal dashed black line indicates a reduction of 99% (MDK_99_) of the surviving population compared to timepoint 0. (B) MDK99 values were interpolated from the killing curves in panel A. Data display five biological replicates and are presented as mean ± SD. For each timepoint, the mutant strains were compared to the WT and statistical significance was assessed by unpaired Student’s t-test. *p < 0.05, **p < 0.01, ***p<0.001

**Table 1:**
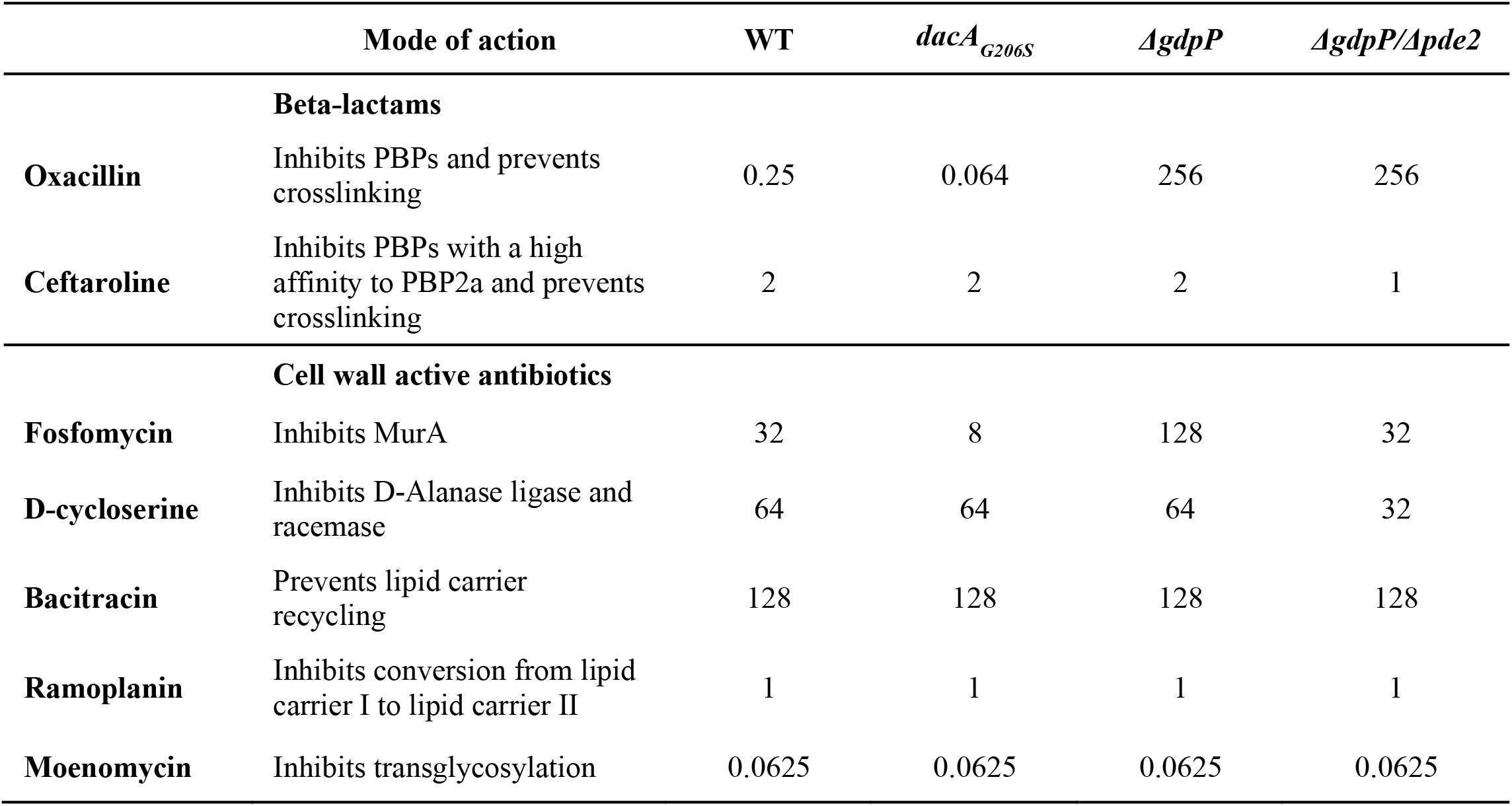
Minimal inhibitory concentrations in [µg/ml] of various antibiotics blocking different steps in the cell wall synthesis.

|  | **Mode of action** | **WT** | *dacA<sub>G206S</sub>* | *ΔgdpP* | *ΔgdpP/Δpde2* |
| --- | --- | --- | --- | --- | --- |
|  | **Beta-lactams** |  |  |  |  |
| **Oxacillin** | Inhibits PBPs and prevents crosslinking | 0.25 | 0.064 | 256 | 256 |
| **Ceftaroline** | Inhibits PBPs with a high affinity to PBP2a and prevents crosslinking | 2 | 2 | 2 | 1 |
|  | **Cell wall active antibiotics** |  |  |  |  |
| **Fosfomycin** | Inhibits MurA | 32 | 8 | 128 | 32 |
| **D-cycloserine** | Inhibits D-Alanase ligase and racemase | 64 | 64 | 64 | 32 |
| **Bacitracin** | Prevents lipid carrier recycling | 128 | 128 | 128 | 128 |
| **Ramoplanin** | Inhibits conversion from lipid carrier I to lipid carrier II | 1 | 1 | 1 | 1 |
| **Moenomycin** | Inhibits transglycosylation | 0.0625 | 0.0625 | 0.0625 | 0.0625 |

Next, we tested whether the susceptibility to cell wall active antibiotics other than beta-lactams was also influenced by c-di-AMP levels. We assessed the MICs of fosfomycin, D-cycloserine, ramoplanin, bacitracin and moenomycin, which all interfere with different steps in the cell wall synthesis (Table 1). Altered c-di-AMP levels affected susceptibility to fosfomycin, which inhibits one of the first steps of the cell wall synthesis. Susceptibility of antibiotics inhibiting later steps in cell wall synthesis were not affected (Table 1). Fosfomycin MICs increased in the Δ*gdpP* mutant and decreased in the *dacA_G206S_* strain (Table 1, lower part), while the Δ*gdpP/*Δ*pde2* mutant displayed unchanged susceptibility to fosfomycin and a slightly increased susceptibility to D-cycloserine as compared to the other strains.

### Mutants with high c-di-AMP levels display a thickened cell wall

The involvement of c-di-AMP in resistance to beta-lactams and fosfomycin as well as the increased peptidoglycan crosslinking found in a *S. aureus* Δ*gdpP* strain (3) raises the question whether cell wall thickness is affected by c-di-AMP levels. Hence, we measured cell wall thickness using transmission electron microscopy (TEM). The Δ*gdpP* and Δ*gdpP*/Δ*pde2* mutants displayed a significantly thicker cell wall than the WT (27.3 ± 4.6 nm, 26.1 ± 4.1nm and 21.0 ± 2.6 nm, respectively) (Figure 4A, B). Lower amounts of c-di-AMP in the *dacA_G206S_* mutant did not affect cell wall thickness (20.2 ± 3.0 nm).

**Figure 4:**
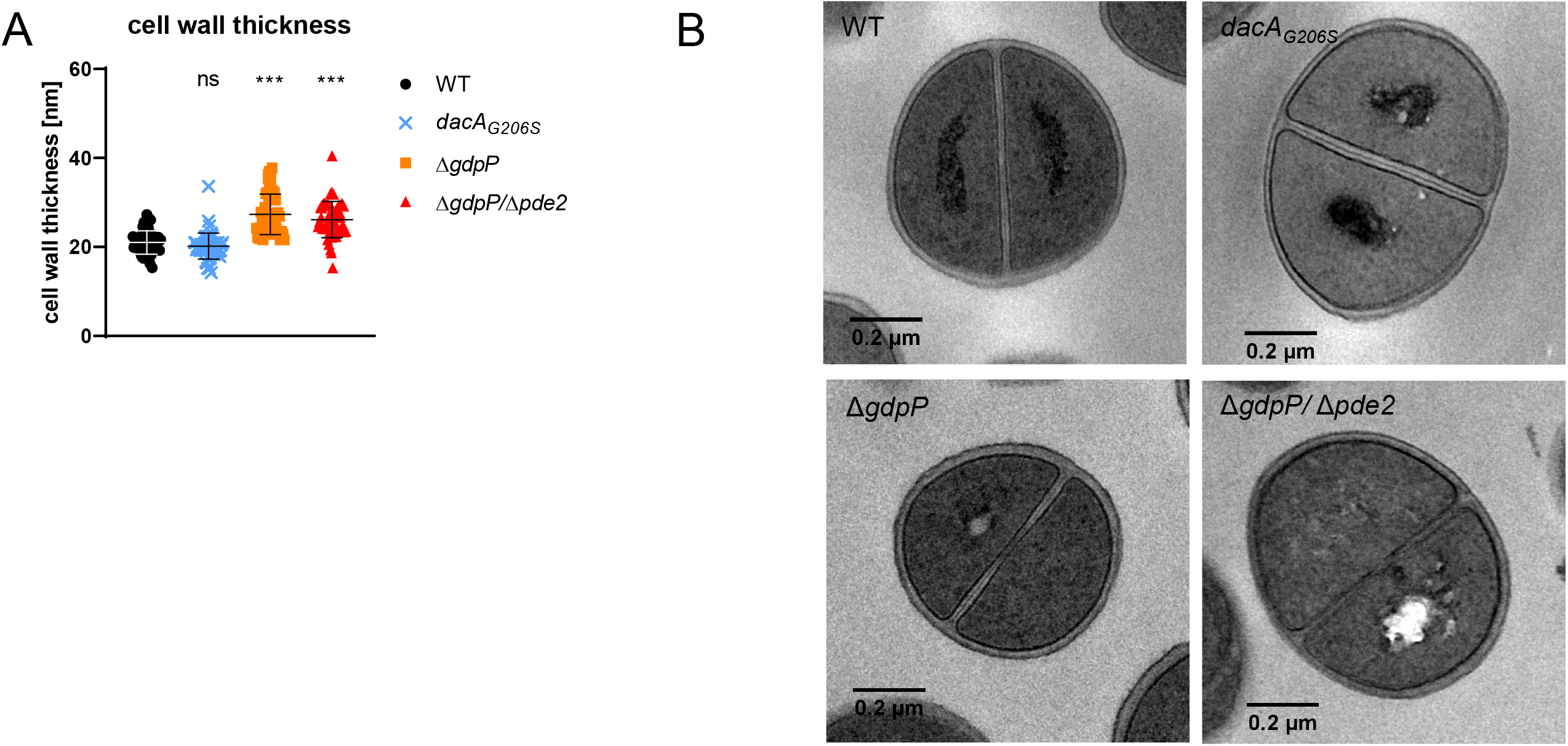
Mutants with high c-di-AMP levels display a thickened cell wall. (A) Cell wall thickness was determined with TEM of bacteria grown for 18 hours in liquid TSB. 50 - 70 cells per strain were analysed with ImageJ. Representative images of each strain are shown in (B). Data display mean ± SD. Each dot represents a single cell (median of five measurements per cell). Statistical significance was assessed by unpaired t-test comparing WT to each strain. *p < 0.05, **p < 0.01, ***p<0.001

### High c-di-AMP levels increase the activation of the cell wall stress stimulon CWSS

Thickening of the cell wall can be caused by a misbalance between autolysis and cell wall synthesis. In *S. aureus*, cell wall synthesis increases in response to antibiotic-induced cell wall stress through differential expression of a set of over 40 genes (13). The activation of this cell wall stress stimulon (CWSS) is controlled by the three-component system VraTSR (22, 23). A link between c-di-AMP levels and CWSS expression has been shown before in a *S. aureus dacA_G206S_* strain (11). We hypothesized that increased c-di-AMP levels can lead to cell wall thickness increase and beta-lactam resistance through activation of the CWSS and tested this hypothesis using the Δ*gdpP* mutant. CWSS basal expression was assessed in the absence of antibiotic-induced stress by quantifying the activity of the *sas016* gene promoter, widely used as a proxy for CWSS activation (11, 24, 25). Confirming our hypothesis, the CWSS activity was increased in the Δ*gdpP* mutant (up to 4.5 times) as compared to the WT (Figure 5A). To investigate whether c-di-AMP regulates CWSS expression through VraTSR signaling, we created a *ΔvraR* single and a Δ*gdpP/*Δ*vraR* double mutant. Deletion of *vraR* reduced CWSS activation to a minimum while a double deletion of *vraR* and *gdpP* only reduced CWSS activation as compared to the Δ*gdpP* mutant, indicating that c-di-AMP can activate CWSS in a VraR- independent manner (Figure 5A). We then investigated whether VraR-dependent CWSS activation influences c-di-AMP levels. Deletion of *vraR* significantly lowered c-di-AMP concentration when compared to the WT strain and a similar albeit not significant trend was found for the Δ*gdpP/*Δ*vraR* mutant, likely due to increased variance of the Δ*gdpP* background, when compared to the Δ*gdpP* mutant (Figure 5B). C-di-AMP levels in the Δ*gdpP/*Δ*vraR* mutant remained significantly above WT level.

**Figure 5:**
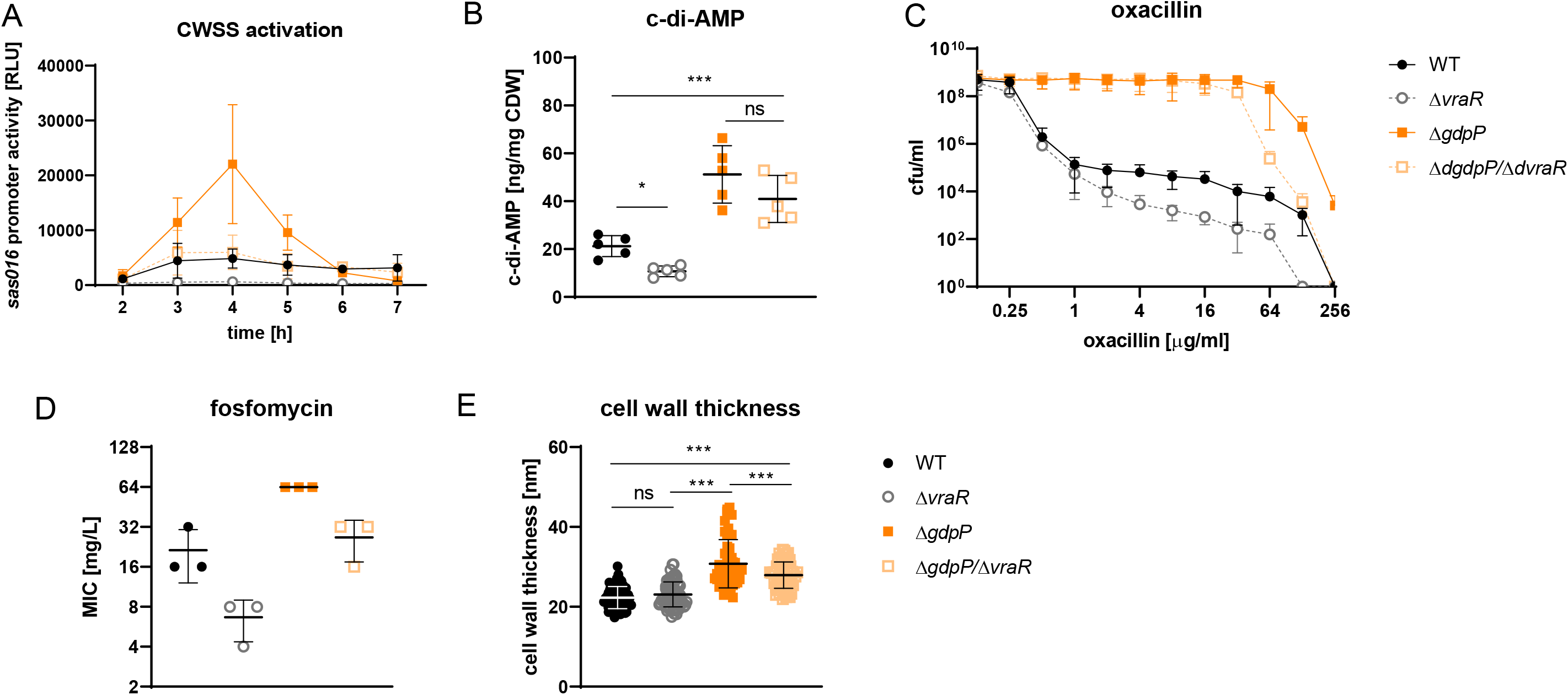
High c-di-AMP levels increase the activation of the cell wall stress stimulon. (A) CWSS activation over 7 hours of growth in TSB containing tetracyclin was measured via the VraR-responsive promotor of the gene *sas016* fused to a luciferase gene (Dengler et al 2013&2016). (B) C-di-AMP was quantified in late exponential growth cultures (OD 2) with LC-MS and normalized to the cellular dry weight (CDW). (C) Resistance to oxacillin is shown in a population analysis profile. (D) MIC of fosfomycin in TSB was measured by broth microdilution. (E) Cell wall thickness was determined with TEM of bacteria grown for 18 hours in liquid TSB. 50 - 70 cells per strain were analysed with ImageJ. Each dot represents a single cell (median of five measurements per cell). Data display mean ± SD of three (A, C, D) or five (B) biological replicates. (B, E) Statistical significance was assessed by one-way ANOVA correcting with Dunnett’s test for multiple comparisons. *p < 0.05, **p < 0.01, ***p<0.001

Increased CWSS activation confers resistance to cell wall active antibiotics (22). The elevated basal CWSS activation in the Δ*gdpP* mutant caused by high c-di-AMP levels might contribute to the resistance to oxacillin and fosfomycin. Deletion of *vraR*, and hence a reduced CWSS activation, should therefore lead to a decrease in resistance. Confirming this hypothesis, we found a decrease in resistance in the Δ*gdpP/*Δ*vraR* mutant as compared to the Δ*gdpP* mutant (Figure 5C, D). However, the double mutant remained more resistant than the WT despite a comparable basal CWSS activation level. The resistance to both antibiotics decreased in the Δ*vraR* single mutant as compared to the WT (Figure 5C, D), in line with earlier studies (25, 26).

Finally, we investigated whether the higher basal CWSS expression level accounted for the thicker cell wall found in the Δ*gdpP* mutant. The cell wall thickness decreased significantly in the Δ*gdpP*/Δ*vraR* double mutant as compared to the Δ*gdpP* mutant, 27.9 ± 3.3 nm and 30.8 ± 6.0 nm, respectively (Figure 5E). However, the double mutant still displayed a thicker cell wall than the WT (22.3 ± 2.8 nm) and the Δ*vraR* strain (23.1 ± 3.1 nm).

In conclusion, we observed increased CWSS activation in the Δ*gdpP* strain, which partially but not completely accounted for thickening of the cell wall and the increase in oxacillin as well as fosfomycin MIC.

## Discussion

In this study, we show that c-di-AMP levels affect *S. aureus* growth, virulence and antibiotic tolerance and resistance. Antibiotic resistance and tolerance are pressing health-care related problems rendering effective treatment of bacterial infections increasingly difficult. Mutations in the c-di-AMP degrading phosphodiesterase GdpP have recently been found to cause resistance to beta-lactam antibiotics in clinical *S. aureus* isolates (8).

In *S. aureus*, *Streptococcus suis* and *Streptococcus pneumoniae*, mutations in either one or both phosphodiesterases degrading c-di-AMP lead to a decreased growth rate (4, 15, 27, 28). Consistent with these findings, we measured clearly reduced growth based on OD measurements and decreased colony size on agar plates in the two mutants characterized by high c-di-AMP levels. However, we could not confirm the growth defect by CFU counts, suggesting variations in cell size as a confounding factor. Our data showed that *S. aureus* cell size negatively correlated with c-di-AMP levels, leading to a substantial reduction in cell volume in the Δ*gdpP* and Δ*gdpP*/Δ*pde2* mutants, characterized by high c-di-AMP levels, and to an increase in cell volume in the *dacA_G206S_* mutant, characterized by low c-di-AMP levels. Single-cell time-lapse microscopy revealed that the increased c-di-AMP level in the Δ*gdpP* mutant did not impact its generation time indicating that the smaller cell volume is responsible for the reduced growth detected by OD measurements. These findings are in line with a recent study showing an increase in cell size in *S. aureus* JE2 correlated with an increase in OD (29). The Δ*gdpP*/Δ*pde2* and *dacA_G206S_*mutants showed a longer generation time indicating a c-di-AMP optimum concentration range for unimpaired growth. Similar results have been found in *Streptococcus pyogenes* where a Δ*gdpP*/Δ*pde2* and a *dacA_G206S_* mutant, but not the Δ*gdpP* single mutant, showed an increased doubling time in liquid medium during exponential growth (30).

Although mutations leading to increased c-di-AMP levels have been found to be relevant for beta-lactam resistance in clinical isolates, it is still unclear how c-di-AMP affects the virulence of these isolates. The antioxidant staphyloxanthin is an important virulence factor that protects *S. aureus* from oxidative stress, encountered during infections in the host environment. We quantified lower levels of staphyloxanthin in the two strains with high c-di-AMP levels. Matching the decreased staphyloxanthin concentrations, survival after exposure to H_2_O_2_ was reduced in strains with increased c-di-AMP levels. This confirms previous findings showing a downregulation of the superoxide dismutase *sodM* in a *S. aureus ΔgdpP* mutant, explaining the decreased ability to cope with oxidative stress (15).

The exact mechanism how c-di-AMP mediates resistance to beta-lactams is not yet known. In general, resistance to beta-lactams in *S. aureus* is mediated by PBP2a encoded by the *mecA* gene (1) or by PBP4 in *mecA*-negative strains (31, 32). Increased *pbp4* expression was found in a *S. aureus ΔgdpP* strain with increased beta-lactam resistance (15). In contrast, a recent study reported unchanged *pbp4* expression in beta-lactam resistant clinical *S. aureus* isolates lacking *mec*A (8). However, the majority of these *mecA*-negative clinical isolates had acquired mutations in *gdpP,* indicating a clinical relevance of strains with increased c-di-AMP levels. Confirming earlier studies, we found drastically increased MICs for the beta-lactam oxacillin in the *ΔgdpP* and *ΔgdpP/*Δpde2 mutants and a decreased MIC in the *dacA_G206S_*mutant (3, 8, 9, 29, 33). In clinics, cephalosporins are often used to treat MRSA infections as they exhibit high affinity to the MRSA-specific PBP2a. MICs to ceftaroline, a fifth-generation cephalosporin preventing peptidoglycan crosslinking, remained unchanged in the *ΔgdpP* and *ΔgdpP/*Δpde2 mutants, suggesting that increased peptidoglycan crosslinking does not affect resistance to this antibiotic (3). This is supported by Ba et al. showing increased resistance to oxacillin but not to cefoxitin, a second-generation cephalosporin, in clinical *mecA*-negative MRSA isolates harbouring a *gdpP* mutation (9). Similarly, unchanged susceptibility to ceftaroline was found in a *S. aureus* strain with a mutated *clp*, a protease involved in various cell wall processes such as peptidoglycan crosslinking, beta-lactam resistance and cell size, leading to a Δ*gdpP*-like phenotype (34). However, despite having similar MICs, time-kill curves showed prolonged survival and an increased MDK_99_ for the Δ*gdpP* and Δ*gdpP/*Δ*pde2* mutants. As antibiotic tolerance is defined by an increase in MDK_99_ (35), high c-di-AMP levels seem to promote tolerance to ceftaroline. This is in line with a study showing that certain mutations in the staphylococcal *gdpP* gene lead to beta-lactam and glycopeptide antibiotics tolerance (33). Not all *gdpP* mutations found in clinical isolates conferred beta-lactam resistance, therefore the question remains whether those mutations might cause tolerance and thereby pave the way to resistance development (8, 10). A role of c-di-AMP in multi-drug tolerance evolution has recently been proposed in *Mycobacterium smegmatis* (36). Our data suggest that increased c-di-AMP levels delay the ceftaroline effect by promoting tolerance, underlining the difficulties in eradicating these bacteria with certain antibiotics.

While the influence of c-di-AMP on the resistance to beta-lactams has been described, data about resistance to other cell wall active antibiotics is still scarce. We tested five antibiotics (fosfomycin, D-cycloserin, bacitracin, ramoplanin and moenomycin) of different classes inhibiting distinct cell wall synthesis steps. Only resistance to fosfomycin differed between the strains, increasing in the *ΔgdpP* mutant and decreasing in the *dacA_G206S_* mutant. Since fosfomycin inhibits MurA, an enzyme catalyzing the formation of the peptidoglycan precursor UDP-MurNAc (37, 38), these observations suggest that c-di-AMP affects the first steps of cell wall synthesis. However, the fosfomycin MIC remained unchanged in the Δ*gdpP*/Δ*pde2* double mutant compared to the WT. A study in *S. pyogenes* revealed distinct Δ*gdpP* and Δ*pde2* phenotypes regarding growth and virulence, suggesting different cellular roles of the two phosphodiesterases (30). In contrast to GdpP, Pde2 mainly degrades pApA, the product of c-di-AMP degradation by GdpP. In a previous study, an accumulation of pApA was found in a *S. aureus* Δ*gdpP*/Δ*pde2* mutant, and it has been suggested that pApA acts as a nanoRNA source and thereby influences the priming of transcription and ultimately gene expression (4, 30, 39).

Resistance to cell wall active antibiotics often correlates with a thicker cell wall in *S. aureus* (31, 34, 40) and c-di-AMP was shown to be involved in cell wall homeostasis (30, 41). We hypothesized that the increased antibiotic resistance and tolerance found in the mutants characterized by high c-di-AMP levels correlate with a thicker cell wall. Confirming our hypothesis, we detected an increase in cell wall thickness in the *ΔgdpP* and *ΔgdpP/*Δ*pde2* mutants. Whether the thicker cell wall is directly responsible for the increased antibiotic resistance and tolerance still needs to be elucidated.

Cell wall thickness is influenced by autolysis, peptidoglycan crosslinking and cell wall synthesis (42, 43). In *S. aureus*, cell wall synthesis can increase upon cell wall stress induced by antibiotics. Such stress triggers the three-component regulatory system VraTSR that regulates the expression of a cluster of more than 40 genes involved in cell wall synthesis, known as cell wall stress stimulon (CWSS) (12, 44). The exact signal leading to the VraTSR cascade is still unclear (45). It was shown that reduced c-di-AMP levels correlate with a decreased basal CWSS activation in two different *S. aureus* strains carrying the *dacA_G206S_* mutation (11). We therefore hypothesized that high c-di-AMP levels could trigger CWSS activation and therefore lead to an increase in cell wall thickness. In line with our hypothesis, we detected increased CWSS activation in the Δ*gdpP* mutant. In the Δ*vraR* single mutant, CWSS activation was abolished, confirming an earlier study (22); however deletion of *vraR* in the Δ*gdpP* mutant reduced CWSS activation only to WT level. This observation suggests that c-di-AMP activates CWSS in a VraR-independent manner. The increased CWSS activation triggered by c-di-AMP might contribute to a thicker cell wall and increased resistance to oxacillin and fosfomycin as the resistance and cell wall thickness phenotype of the Δ*gdpP/*Δ*vraR* double mutant lies between the WT and the Δ*gdpP* single mutant. Additionally, loss of *vraR* lowered c-di-AMP levels in the WT strain and the Δ*gdpP* mutant, indicating a role of *vraR* in c-di-AMP homeostasis. An opposite effect was found in *Enterococcus faecalis* where a *liaR*, which is a *vraR* homologue, mutant was characterized by increased c-di-AMP levels without changes in the expression of the enterococcal *dacA* and *gdp*P homologues (46), suggesting a species-specific interaction between c-di-AMP levels and the VraTSR regulatory system.

In conclusion, our study shows that a previously described growth defect in *S. aureus* producing high c-di-AMP levels is mainly attributable to smaller cell size. High levels of c-di-AMP impaired virulence by reducing resistance to oxidative stress and staphyloxanthin production, resulted in a thicker cell wall promoting increased resistance to oxacillin and fosfomycin as well as tolerance to ceftaroline and lead to higher CWSS activation. Taken together, we report the importance of studying the role of c-di-AMP in promoting antibiotic tolerance and increasing cell wall thickness, potential mechanisms leading to the development of antibiotic resistance, with the aim of streamlining antibiotic treatment and ultimately improve patient care.

## Materials and Methods

### Bacterial strains and growth conditions

Bacterial strains and plasmids used in this study are listed in Supplementary Table 2. Strains were grown in TSB (tryptic soy broth, BD) at 37°C shaking or on TSB agar plates. Tetracycline was used at 10 µg/ml, chloramphenicol at 10 µg/ml, anhydrotetracycline at 0.2 µg/ml when appropriate. Overnight cultures were grown for 14 to 18 hours, diluted 1:50 in fresh medium and grown for 1.5 hours to obtain logarithmically growing bacteria.

### Plasmid transduction and genetic manipulations

The VraR deletion mutant was constructed by inserting two stop codons by homologous recombination at the second and third codon of the *vraR* gene using a pKOR1 plasmid (25). pKOR1-VraR::stop was transduced into the LAC* WT and the Δ*gdpP* mutant background using phage 80α and chromosomally integrated as previously described (47). LAC* *dacA_G206S_* was constructed using the pKOR1-*dacA-*SNP plasmid from Dengler et al. 2013 (11). The plasmid was transduced into LAC* WT using phage 80α and chromosomally integrated. The manipulated regions were checked by Sanger sequencing. The *sas016* promoter luciferase fusion plasmid p*sas016*_p_-*luc* (*24*)was transduced into LAC* WT, Δ*gdpP* and Δ*gdpP/*Δ*vraR* mutant background using phage 80α.

### Colony growth dynamics quantification

Bacteria from stationary or logarithmic growth phase with or without exposure to H_2_O_2_ stress were diluted to reach 50 to 150 CFUs per COS plate. Plates were incubated at 37 °C and images were acquired with a Canon EOS 1200D reflex cameras every 10 min for 48 h. Cameras were triggered by Arduino Uno board and optocouplers. Colony growth dynamics were analyzed as previously described using ColTapp (48).

### Determination of cell volume and generation time using single cell microfluidics

The size and generation time (time from birth to division of a given cell) of single bacterial cells was determined using a mother machine microfluidics device combined with timelapse microscopy. Device design and fabrication has been described elsewhere (49). In brief, cells were embedded in chambers of height 0.93 µm, length 25 µm, and width ranging from 1.2 to 1.6 µm perpendicular to a flow channel. Bacteria were grown overnight in TSB supplemented with 0.01% Tween20 to avoid clumping. The cultures were diluted 1:100, regrown for two hours and then washed in TSB 0.1% Tween20 and concentrated in TSB 0.01% Tween20. Bacteria were then loaded into the microfluidics chip with a pipette. The outlet and inlets were connected with tubings (Adtech, PTFE, inner diameter: 0.3 mm, outer diameter: 0.76 mm) to a waste collection container and to a syringe pump (KF Technology, NE-1600) providing fresh TSB 0.01% Tween20 at a flow rate of 0.5 ml/h, respectively. The chip was placed inside the heated chamber (37°C) of the microscope into fully automated inverted microscope (Olympus IX83 P2ZF) controlled via CellSense 3.2. After letting the bacteria recover for four to five hours from the loading procedure, imaging was started. Phase contrast (200 ms exposure, 130 intensity IX3 LED) images were acquired every 2 min for 5 h with a 100×, numerical aperture 1.3 oil phase objective (Olympus) and an ORCA-flash 4.0 sCMOS camera (Hamamatsu). A custom MATLAB (MathWorks) script was used to register image sequences, detect filled chambers and crop out single chamber images. On these, single cell segmentation was performed with a retrained deep-learning StarDist 2D model (50). The segmentations were then injected in the retrained deep-learning cell-tracking model of DeLTA 2.0 (51). DeLTA 2.0 output was filtered to include only cells for which we observed a mother cell and a division event. We reported time between division events as generation time and mean cell volume over the lifetime of a cell. Cell volume was estimated as described before (52) using equation 1 (with a and b corresponding to the longer and shorter semi-axes, respectively).

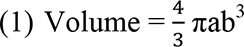

### Susceptibility to H_2_O_2_

Stationary growth phase bacteria were diluted to approximately 5*10^6^ CFU/ml in PBS and incubated for one hour at 37°C with 30 mM H_2_O_2_. Bacterial survival relative to the control (no H_2_O_2_) was determined by plating serial dilutions on blood plates (Columbia agar plates with 5% sheep blood, COS, Biomérieux) in order to inactivate any residual H_2_O_2_.

### Staphyloxanthin production

Staphyloxanthin production was quantified as described before (53) with a few modifications. In short, 2 ml of a stationary growth phase culture (24 h, TSB) were harvested by centrifugation (maximum speed, 5 min, room temperature) and washed with PBS. The washed pellet was completely dried with a centrifuge concentrator (Eppendorf) for 1.5 hours at 30°C. After vortexing the pellet in 20 µl PBS containing glass beads, 180 µl of methanol were added and the samples were kept shaking (1000 rpm) at 55°C for 4 minutes. After centrifugation (maximum speed, 10 min), the absorbance of the supernatant was measured with a plate reader (Tecan Infinite M Nano) at OD_465_.

### Transmission electron microscopy (TEM)

Overnight bacterial cultures grown in TSB were washed in PBS and resuspended in cacodylate Buffer (0.1 M) with 2.5% glutaraldehyde. The samples were processed for image analysis as previously described (54). Images were taken with a 120-kV transmission electron microscope (FEI Tecnai G2 Spirit) equipped with two digital CCD cameras. Cell wall thickness was assessed with ImageJ in 50-70 cells per strain by manually measuring the observed cell wall in zones that are not in proximity of the division septum. The median of five measurements per cell was used to calculate the mean cell wall thickness of each strain.

### Minimal inhibitory concentrations (MICs)

MICs were determined in TSB with the broth microdilution method according to EUCAST guidelines (55). Oxacillin MICs were assessed using commercial MIC test strips (Etest, Biomérieux). MICs were assessed in biological triplicates.

### Oxacillin population analysis profiles

Antibiotic resistance population profiles were determined by plating appropriate dilutions of an overnight culture, ranging from undiluted to 10^-7^, on increasing concentrations of oxacillin. Plates were incubated at 37°C and colony forming units per ml (CFU/ml) were determined after 48 hours.

### Time-kill curves

Bacteria were grown to logarithmic growth phase as described above, washed and inoculated in TSB containing ceftaroline at a concentration of 40x MIC (inoculum 5*10^5^ cfu/ml). After 1.5, 3, 6 and 24 hours, bacterial numbers were assessed by plating and the survival fraction was calculated relative to the inoculum. The values for the minimal duration time to kill 99% of the population (MDK_99_) was interpolated using linear regression.

### Luciferase reporter assay

To measure luciferase activity over different growth stages, cultures were inoculated at OD_600_ 0.05 and samples were collected after 2, 3, 4, 5, 6 and 7 hours of growth and pellets were stored at −20°C until further use. Pellets were defrosted and resuspended to OD_600_ 10 in PBS. Immediately after the addition of the luciferase substrate (Promega) at a 1:1 ratio, luminescence was measured with a plate reader (Spectramax i3, Molecular Devices) using the standard settings for luminescence measurements.

### C-di-AMP quantification

C-di-AMP was quantified as described previously with some modifications (11). Briefly, bacteria were grown to log phase (Supplementary Table 1) or to approximately OD_600_ 2 (Figure 5), collected by fast filtration and immediately quenched (−20°C cold acetonitrile 60%, formic acid 0.5 M 20%, methanol 20%). C-di-GMP (Biolog) was added as an internal standard. Samples were sonicated four times for 20 s before being snap frozen in liquid nitrogen and lyophilized. Analysis was performed on a Thermo Ultimate 3000 UHPLC system (Thermo Scientific, CA, USA) coupled to Thermo OExactice plus instrument (Thermo Fisher Scientific, Waltham, MA, USA) equipped with a heated electrospray ionization probe. LC separation was carried out at a flow rate of 500 µl/min applying a quaternary gradient, summarized as binary gradient. Solvent A was 10mM ammonium formate adjusted to pH 8.1 and solvent B was 10mM ammonium formate adjusted to pH 8.1 in methanol to water in a ratio of 80:20. Gradient B was as follows: 0 min, 2.5%; 4.2 min, 28.8%; 8.9 min, 97.5%; 9.9min, 97.5%; 10.0 min, 2.5%; 13.1 min, 2.5%. MS spectra were acquired in the positive FTMS mode at mass resolution of 70000 (m/z = 200). Five biological replicates per strain were analyzed. For normalization (Supplementary Table 1), total protein content was assessed with the Pierce BCA protein assay kit (Thermo Scientific) according to the manufacturer’s instructions. C-di-AMP concentration was normalized to the internal standard (A_internal_) and OD/total protein ratio (OD_perTP_) using equation (2).

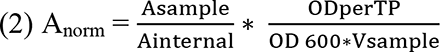

C-di-AMP concentration of the *vraR* mutant strains (Figure 5) were normalized to the cellular dry weight as described before (11).

### Statistical analysis

Data was analyzed with GraphPad Prism 8.0, R 4.3 and RStudio 1.3. Cell volume, generation time and colony size were assessed with a mixed effects model using a random intercept term for the biological replicate followed by estimated marginal means *post-hoc* tests (multivariate t-distribution based p-value correction) (56). All other experiments were analyzed with unpaired Student’s t-tests comparing the WT to each other strain.

## Acknowledgments

We thank Angelika Gründling for providing the *S. aureus* Lac* WT, Δ*gdpP* and Δ*gdpP/pde2* strains.

