## supplementary figure and table for "C-di-AMP levels modulate *Staphylococcus aureus* cell wall thickness as well as virulence and contribute to antibiotic resistance and tolerance"

### Supplementary Table S1

| strain | c-di-AMP (log2 fold change rel. to WT) | p-value |
| --- | --- | --- |
| *S. aureus* LAC* *dacA*$_{G206S}$ | -2.58 ± 0.56 | 7.19e-05 |
| *S. aureus* LAC* *ΔgdpP* | 1.03 ± 0.46 | 1.56e-03 |
| *S. aureus* LAC* *ΔgdpP/Δpde2* | 1.61 ± 0.47 | 1.17e-04 |

**C-di-AMP levels.** C-di-AMP levels of bacteria grown to exponential growth phase (1:50 dilution of overnight cultures grown for 1.5 hours) in TSB was measured by LC-MS and normalized to the total cellular protein content. The fold change (log2 transformed) relative to the WT strain was calculated. Data display mean ± SD of five biological replicates, p-value was calculated by unpaired Student's t-test comparing WT to each strain including error propagation of the total protein measurement.

### Supplementary Table S2

| strain | description | reference |
|---|---|---|
| *S. aureus* LAC* WT | LAC*, erythromycin-sensitive CA-MRSA LAC strain cured of pUSA03 | Corrigan et al. 2011 |
| *S. aureus* LAC* $dacA_{G206S}$ | LAC* *dacA206S* containing a *dacA* with a SNP resulting in a Gly206Ser substitution | This study |
| *S. aureus* LAC* *ΔgdpP* | LAC*Δ*gdpP*::*kan* | Corrigan et al. 2011 |
| *S. aureus* LAC* *ΔgdpP/Δpde2* | LAC* Δ*gdpP*::*kan* Δ*pde2*::*erm, gdpP/pde2* | Bowman et al. 2016 |
| *S. aureus* LAC* *ΔvraR* | LAC* containing *vraR* mutation, truncating VraR after the 2nd amino acid | This study |
| *S. aureus* LAC* *ΔgdpP /ΔvraR* | LAC*Δ*gdpP*::kan containing *vraR* mutation, truncating VraR after the 2nd amino acid | This study |
| **plasmids** | | |
| pKOR1-VraR::stop | pKOR1 construct containing the mutated *vraR* gene with two in-frame stop codons inserted between the 2nd and 3rd *vraR* codons | McCallum et al. 2011 |
| pKOR1-*dacA*-SNP | pKOR1 construct containing the *dacA* gene with 1000-bp upstream and 1020-bp downstream regions carrying a mutation leading to Gly206Ser substitution in DacA | Dengler et al. 2013 |
| p*sas016*p-luc+ | pBUS1 containing the *sas016* promoter-luciferase reporter gene fusion | McCallum et al. 2011, Dengler et al. 2016 |

### Supplementary Figure S1

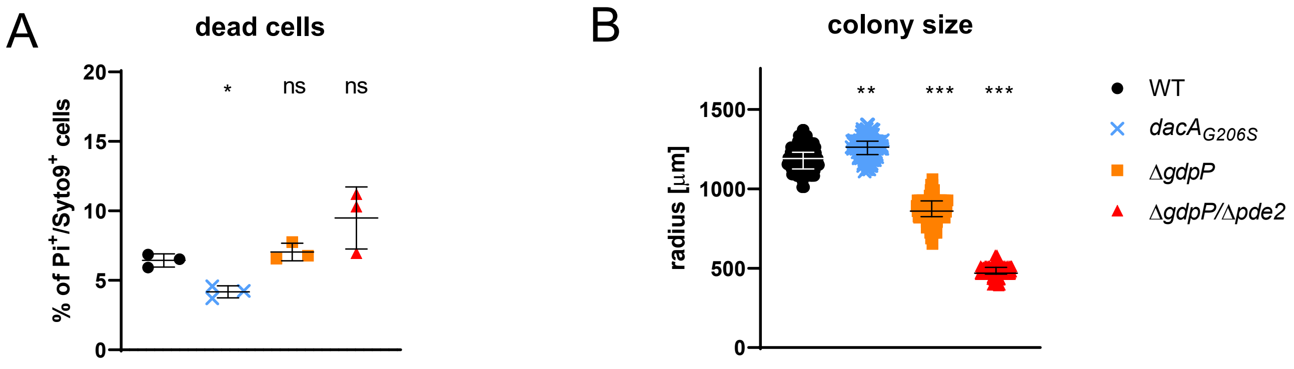


**Figure S1: Percentage of dead cells in logarithmic growth phase.** (A) Bacteria were grown to exponential growth phase (1:50 dilution of overnight cultures grown for 1.5 hours) in TSB, washed and stained with Syto9 (all cells) and PI (dead cells). Fluorescence was analyzed with a flow cytometer and the percentage of PI+/Syto9+ cells was calculated. (B) Colony size was assessed on agar plates (50-150 colonies per plate) with a time-lapse imaging approach. Data displays (A) mean +/- SD or (B) median and interquartile range of three biological replicates with each dot representing one colony. Statistical significance was assessed by unpaired Student's t-test comparing WT to each strain on the (A) mean or (B) median values of the biological replicates. *$p < 0.05$, **$p < 0.01$, ***$p<0.001$

### Supplementary Movie S1

**Movie S1: Representative movies of microfluidic experiments.** A: WT, B: *dacA*$_{G206S}$, C: Δ*gdpP* and D: ∆*gdpP*/∆*pde2.* Movies are the basis for the analaysis of the cell volume and generation time shown in Figure 1C and D. Experimental and imaging details are described in the material and methods section.

#### Supplemental methods

##### Detection of live and dead bacteria by flow cytometry

Logarithmic growth phase cultures were diluted to $OD_{600}$ of 0.5 in PBS. For the identification of live and dead bacteria, staining was performed with Syto 9 and propidium iodide (PI, both Molecular Probes). In short, 200 µl of bacterial suspension were mixed with 6 µl of Syto9 and PI each (final concentrations 0.15 µM and 1.32 µM, respectively) and incubated for 2 minutes in the dark at room temperature. Samples were analyzed with Attune NxT flow-cytometer (ThermoFisher) with SSC at 280 and FSC at 100. The population was gated first for bacteria and then single cells. The single cell gate was used for the analysis of PI- and Syto9 double-positive cells (dead bacteria) using the BL1 and YL2 lasers.
